# Exploring transcriptomic and genomic latent variable correction approaches in differential expression analysis

**DOI:** 10.64898/2026.04.07.716914

**Authors:** Yadusayan Appulingam, John Jammal, Aminah Ali, Simon Topp, NYGC ALS Consortium, Alfredo Iacoangeli, Oliver Pain

## Abstract

**Background:** Differential expression analysis is a central tool for studying the biological processes altered in human diseases via transcriptomic signatures. However, transcriptomic datasets are systematically confounded by latent variables from two distinct sources: unmeasured technical and biological heterogeneity within the expression data, and expression differences driven by population stratification. Correction using expression-based surrogate variables (SVs) and genotype-based principal components (PCs) addresses these sources independently, yet no study has directly evaluated their combined use against either method alone within a differential expression framework. In this study we hypothesised that simultaneously including both correction layers would produce more biologically valid and reproducible results than either approach alone, and tested this in two independent RNA-seq datasets of amyotrophic lateral sclerosis (ALS) cases and controls with matching genotype data.

**Results:** Four nested differential expression models (corrected for PC-only, SV-only, both SV and PC, and neither PCs nor SVs) were evaluated across the KCLBB (96 cases and 52 controls) and ALS Consortium (272 cases and 35 controls) datasets. Models were evaluated on: cross-dataset effect size concordance, cross-dataset replicability quantified by the Jaccard Similarity Index, and biological recall against a curated reference set of 66 known ALS genes. The combined SV+PC framework consistently outperformed simpler models across all metrics. Replicability improved nearly ten-fold compared to the non-corrected model, (Jaccard index: 2.28% to 19.5%), and the combined framework exhibited a statistically significant 2.1% gain over the SV-only model. The biological recall ALS genes recovered doubled comparing to the SV correction alone. Crucially, effect size stability was preserved, with the combined model expanding the shared transcriptomic signal without sacrificing consistency. These findings remained generally robust to PC number in sensitivity analyses.

**Conclusions:** This study found that SVs and genotype PCs address non-redundant sources of confounding, and we recommend their combined use as standard practice in differential expression analysis where matched genotype data are available. Notably PCs capturing population structure can also be derived directly from RNA-seq data, extending the applicability of this framework to studies lacking matched genotype data. Although this analysis was restricted to ALS datasets, we expect these findings to generalise to other traits.

## Introduction

Complex diseases such as amyotrophic lateral sclerosis (ALS) are characterised by multifactorial aetiologies in which genetic, environmental, and biological factors interact to cause disease [1,2]. The resulting biological and clinical landscapes of ALS are highly heterogenic, and this is reflected at molecular level in its transcriptome [3,4,5]. Understanding the transcriptional landscape of ALS is therefore inherently challenging, and differential expression analysis is a central tool for identifying disease-associated transcriptomic signatures. However, a substantial proportion of variation in any transcriptomic dataset is driven by latent variables, factors that influence observed measurements but are not explicitly captured in study metadata. These include technical sources such as batch effects, library preparation inconsistencies and sequencing platform variation, as well as biological sources such as cell-type composition, tissue heterogeneity and genetic background [6]. Because latent variables simultaneously affect the expression of many genes, they introduce spurious associations, reduce statistical power, and generate irreproducible results [6,7]. The failure to account for this hidden variation risks producing findings that reflects the idiosyncrasies of a particular dataset rather than true disease biology.

Two distinct types of latent variation are particularly relevant in transcriptomics. The first arises from unmeasured technical and biological confounders within the expression data itself. Surrogate variable analysis (SVA) addresses this by decomposing the residual expression matrix to identify latent factors that explain more variation than expected by chance, constructing surrogate variables (SVs) that are then included as covariates in the differential expression model [6]. The second arises from population stratification, which refers to broader systematic differences across ancestrally distinct subgroups. These manifest in gene expression patterns, partly as expression is heritable and regulated by common genetic variation [8], but also through other ancestry-related biological and environmental factors. Principal component analysis (PCA) of genome-wide genotype data produces genotype-derived principal components (PCs) that can be included as covariates in regression models to account for genetic latent variation. While PCs can capture multiple sources of systematic variation, including technical artefacts, in practice the leading PCs typically reflect population stratification across samples and are widely used to correct for this [9]. In transcriptomic studies, we therefore assume the primary effect of including genotype PCs reflects their correlation with population structure, though this cannot be confirmed directly, and it is equally possible that SVs partially absorb population-specific variation, though the extent to which this occurs is also poorly characterised. Critically, because SVA operates on the expression data alone and is constrained to protect the primary variable of interest [6], it may not fully remove confounding driven by population stratification, particularly when it is correlated with disease status. Recent evidence suggests that population-specific variation accounts for only a small proportion of total expression variance relative to individual heterogeneity and batch effects [10], meaning SVs are likely driven primarily by larger hidden batch effects, leaving residual expression variation related to population stratification unaddressed. Including genotype PCs is therefore a complementary and potentially non-redundant correction strategy. Importantly, genetic PCs do not require separate genotyping, as they can be derived directly from RNA-seq data itself, which has been shown to be yield results comparable to those from array data [11].

Despite the widespread use of latent variable correction in differential expression analysis, there is a lack of systematic evaluations to determine which correction strategy, or combination of strategies, produces the most biologically valid and reproducible results. The methodological literature has focused predominantly on comparisons between statistical models for count data (e.g. DESeq2, edgeR and limma-voom) rather than on the covariate structure of the model itself [12,13]. In practice, approaches vary considerably. Some studies apply SVA alone to control for hidden confounding [14], others rely on expression PCs [15], and genotype PCs are rarely incorporated into differential expression analysis models [11]. While one study employed both genotype PCs and expression derived SVs simultaneously [16] and another ALS blood transcriptomic study utilised both expression derived and genotype PCs [17], implicitly recognising these as distinct and non-redundant layers of confounding, to our knowledge, no systematic benchmark has directly evaluated SVs alone, PCs alone and a combined SV+PC approach within a single differential expression analysis framework, leaving it unresolved whether both correction layers add meaningful, non-redundant value.

Big consortia, where large-scale datasets are aggregated across multiple clinical sites, represent an example of a scenario where unknown batch effects, variable RNA integrity and diverse population structure introduce effects on gene expression analysis that are challenging to account for. We therefore hypothesise that the simultaneous inclusion of genotype PCs and expression SVs in a differential expression analysis framework will produce more biologically valid and reproducible results than either strategy applied in isolation. To test this, we evaluated four nested models in which differential gene expression analysis is corrected with PC-only, SV-only, SV+PC and neither of them (Naïve), across two independent post-mortem motor cortex RNA-seq ALS datasets, evaluating each model’s effect size consistency, cross-dataset replicability and biological recall of established ALS genes.

## Methods

### Study Design and Datasets

Bulk RNA-seq and whole-genome sequencing (WGS) data were analysed from post-mortem motor cortex tissue across two independent ALS cohorts. To account for case-control imbalances, effective sample size (*N_eff_*) was calculated per cohort using: *N_eff_* = 4/(^1^/*N_cases_* + ^1^/*N_controls_*) [18]. Final cohorts were restricted to samples with overlapping RNA-seq, WGS and clinical data.

The first dataset was sourced from King’s College London and MRC London Neurodegenerative Diseases Brain Bank (KCLBB) [19,20], including 148 samples (96 ALS cases, 52 controls; *N_eff_* = 134.92). The second was from the New York Genome Center (NYGC) ALS Consortium initiative (ALS Consortium), comprising 307 samples (272 ALS cases, 35 controls; *N_eff_* = 124.04), with RNA-seq data collected from the medial and unspecified motor cortex sub-regions.

### Transcriptomic Pre-processing and Quality Control

The same bioinformatics pipeline was applied across both datasets, based on the framework established by Kabiljo et al. [14]. Sex chromosomes were excluded from raw count matrices to prevent sex-linked biases from obscuring population structure and disease signal. Normalisation was performed using the median-of-ratios method implemented in DESeq2 (version 1.48.2) [21] to correct for differences in sequencing depth and library composition. Genes were retained if they reached a minimum normalised count of 5 in at least 10 samples, reducing the multiple-testing burden and removing low abundance transcripts likely representing technical noise rather than true biological signal.

Continuous covariates were assessed for linearity with log2 normalised counts prior to model inclusion. RNA Integrity Number (RIN) was converted to a categorical variable (RIN_Cat_) as it did not maintain a linear relationship with expression. Age at Death and Post-Mortem Delay (PMD) demonstrated sufficient linearity and were included as centred continuous variables. PMD was excluded from the ALS Consortium model due to high missingness. A site covariate was included in the ALS Consortium model to account for multi-site sample collection, a source of systematic technical variation not present in KCLBB.

### Genomic Data Quality Control and PCA

Genotype data was processed using PLINK2 [22], applying standard quality control filters. Variants were retained if they were biallelic, had a minor allele frequency (MAF) > 0.05, genotype missingness < 0.02 and passed Hardy-Weinberg Equilibrium testing (*P* > 1 × 10^+,^). Linkage disequilibrium (LD) pruning was performed (--indep-pairwise 50 5 0.2) and long-range LD regions were manually excluded to prevent them from disproportionately influencing PC estimation [23]. Missing genotypes were mean imputed to facilitate PCA. PCA was performed using the bigsnpr package [24]. For KCLBB, PCs were calculated using all 247,287 variants remaining post-QC. In ALS Consortium, homozygous reference genotypes were encoded as “./.” and were therefore indistinguishable from genuinely missing genotypes. Consequently, applying a standard genotype missingness filter (<0.02) excluded a large proportion of variants, because many sites appeared to have high missingness despite containing true homozygous reference calls. As a result, PCA was effectively restricted to variants with relatively few homozygous reference calls and a higher proportion of heterozygous or homozygous alternate genotypes, yielding 26,906 LD-independent variants. Five genotype-derived PCs were retained per cohort based on visual inspection of scree plots, selecting PCs up to the elbow of the plot beyond which the proportion of variance explained showed negligible incremental change (Supplementary Figure S1).

Global ancestry was inferred for all samples using the GenoPred pipeline, which maps cohort genotypes against a reference panel comprising from 1000 Genomes Project (2015) and the Human Genome Diversity Project [25].

### SVA

Expression-derived SVs were estimated using the Leek method via the sva package in R [26], which determines the number of SVs automatically by a data-driven approach based on the residual expression matrix after regressing out known covariates. One SV was generated for KCLBB and three for ALS Consortium. Prior to modelling, systematic association testing was performed to evaluate independence between SVs, genotype PCs and measured clinical and technical covariates, assessing the risk of multicollinearity in combined models (full details in Supplementary Methods).

### Differential Expression Modelling Framework

Differential expression was performed using DESeq2 [21], which models count data within a negative binomial generalised linear model. The primary coefficient of interest represents the log2 fold change in gene expression associated with ALS disease status. Four nested models were benchmarked to evaluate the incremental impact of latent variable correction.

**1**. **Naïve model**: The baseline model which includes primary clinical and technical covariates.

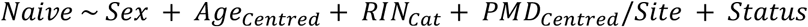
**2**. **PC-only model**:

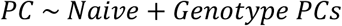

Five genotype PCs were included per model.
**3**. **SV-only model**:

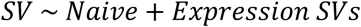

One SV was included for KCLBB and three for ALS Consortium.
**4**. **SV+PC model (Combined)**:

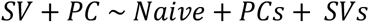

Full model specifications for each cohort are provided in Supplementary Methods. Multiple testing correction was applied using Independent Hypothesis Weighting (IHW), implemented via the IHW package in R [27]. IHW weights genes by mean expression level, improving statistical power over the standard Benjamini-Hochberg procedure while maintaining FDR control at 0.05.

### 2.6 Model Evaluation

*Intra-dataset characterisation.* Within each cohort, the sets of differentially expressed genes (DEGs; IHW-adjusted FDR < 0.05) identified across the four models were visualised using four-way Venn diagrams to quantify unique and shared discoveries. Effect size concordance between models was assessed using Pearson’s (*r*) and Spearman’s (ρ) correlation of log2 fold change (LFC) estimates, evaluated across three gene sets: the intersection of significant DEGs, the union of significant DEGs, and all filtered genes. Pearson’s correlation measured linear consistency in effect size magnitude, while Spearman’s rank correlation provided a non-parametric assessment robust to the influence of outliers.

*Cross-dataset benchmarking of effect size.* To evaluate consistency of LFC concordance for the same model between KCLBB and ALS Consortium, the same comparisons across the intersect of significant DEGs, union of significant DEGs and the shared gene universe was utilised. A higher cross-dataset correlation was interpreted as evidence that a model successfully decoupled the conserved ALS transcriptomic signature from dataset specific confounding. To determine if observed difference in Pearson (*r*) and Spearman (ρ) correlations between SV and SV+PC frameworks was statistically significant, bootstrap resampling (1000 iterations) was performed via the boot package in R [28]. We generated 95% confidence intervals (*CIs*) for Δ*r* and Δρ across the union of significant DEGs, excluding zero to indicate statistical significance.

*Replicability index.* The Jaccard Similarity Index (*J*) was used to quantify the proportion of shared DEGs between cohorts:

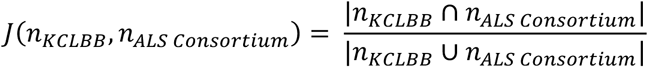

Where n represents the number of significant DEGs per model, ∩ denotes the intersection (DEGs significant in both cohorts) and ∪ represents the union (DEGs significant in at least one cohort). To determine whether the difference in *J* between the SV and SV+PC models (Δ*Jaccard*) was statistically significant rather than attributable to random fluctuation, bootstrap resampling was performed using the boot package in R [28], generating a 95% CI for Δ*Jaccard*, where a CI excluding zero was considered significant. The proportion of overlapping significant DEGs with the same direction of effect was assessed to determine whether differences in replicability were accompanied by consistent directionality of effects across cohorts.

*Biological recall and ‘apparent’ precision.* Model sensitivity was evaluated against a curated reference set of 73 established ALS-associated genes (see Supplementary Table S1). The reference set was derived from the ALS Online Database (ALSoD) [29] and refined through expert curation based on published ALS genetic literature, retaining genes with strong evidence of ALS association from peer-reviewed studies. Seven genes were subsequently excluded following failure to meet expression thresholds in at least one cohort, yielding a final reference set of 66 genes. Herein, we refer to this list of established ALS genes as ‘*known ALS genes*’. Biological recall was defined as:

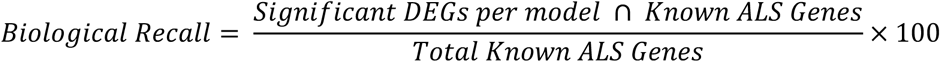

This metric directly tests whether more stringent latent variable correction enriches the DEG list for known disease-relevant biology without sacrificing sensitivity.

To complement recall, ‘apparent’ biological precision was calculated as:

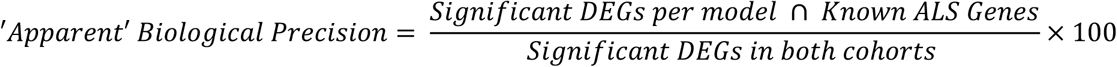

This determines whether the differences in ALS genes is proportionally reflected in the differences in total DEGs. ‘Apparent’ reflects the inherent limitation that known ALS reference is necessarily incomplete.

### Sensitivity Analysis

To evaluate the robustness of our findings to the number of genotype PCs included as covariates, a sensitivity analysis was performed across all three metrics: cross-dataset effect size consistency, Jaccard replicability index and biological recall. The number of PCs included in the SV+PC model was varied independently for both KCLBB and ALS Consortium across intervals of PC1, PCs 1-3, PCs 1-5, PCs 1-7, PCs 1-10, yielding 25 comparisons per metric. For each comparison, performance of the SV+PC model was assessed relative to the SV-only model.

## Results

### Multicollinearity and Covariate Selection

Prior to differential expression analysis, systematic testing showed that most latent variables were not strongly associated with measured covariates, although several notable associations were identified. In KCLBB, SV1 showed a significant association with RIN (p_adj_ = 0.007), though the shared variance was negligible (r = –0.249, r^2^ = 0.062; Supplementary Table S2). In ALS Consortium, SV1 was significantly associated with collection site, suggesting that SVs capture some site-specific variation, though we cannot determine whether it is technical artefacts, population stratification or a combination of both. Several genotype PCs (PC1, PC2 and PC4) also associated with site (most significant p_adj_ < 2.66 x 10^-18^; Supplementary Table S2), consistent with population stratification across geographically distributed sites. However, it is possible that this partly reflects technical batch effects between genotype and RNA-seq data collection procedures. Notably, ALS Consortium PC1 was significantly associated with disease status, indicating that the leading axis of genetic structure in this cohort was not independent of the case-control comparison. PC1 was nevertheless retained in the model because genotype PCs were included to account for broad genetic structure, although we acknowledge that in observational case-control data such adjustment may remove both confounding variation and status-associated biological signal. Critically, no statistically significant associations were identified between SVs and PCs in either cohort following BH correction (maximum r = 0.18 in KCLBB; –0.15 in ALS Consortium; Figure 1), providing no evidence of multicollinearity between genomic and transcriptomic latent factors. All latent variables were therefore retained in the combined SV+PC model, as PCs and SVs represent independent signals extending beyond the variance captured by site or RIN_Cat_ alone.

**Figure 1.**
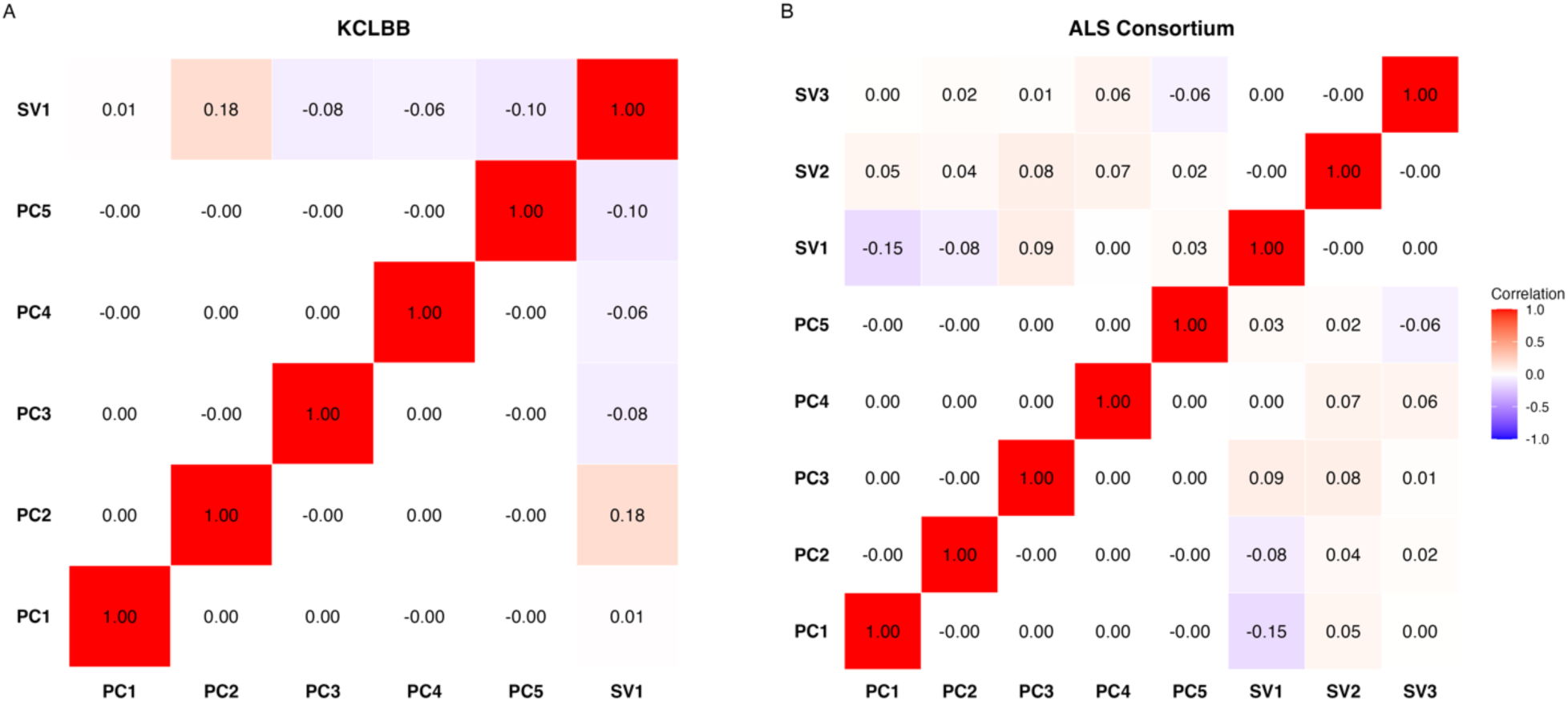
Pairwise correlation matrices of genomic and transcriptomic latent variables. Correlation matrices illustrating the relationship between genotype-derived PCs and expression derived SVs for KCLBB (A) and ALS Consortium (B) cohorts. The colour scale and circle area represent the magnitude of the Pearson’s correlation coefficient (r), ranging from –1 (blue) to +1 (red). The diagonal represents the self-correlation of each variable (r = 1.00), while off-diagonal elements indicate the correlation between distinct latent factors.

### Global Ancestry

Ancestry inference confirmed meaningful differences in cohort composition between the two datasets (Figure 2). KCLBB demonstrated high genetic homogeneity, with all but one sample assigned to European ancestry (posterior probability > 0.95). In contrast, ALS Consortium exhibited greater diversity: 262 European, 7 African, 6 American, 2 Central/South Asian, 3 East Asian and 27 admixed or unassigned samples. The difference in ancestral composition between cohorts provided a valuable opportunity to assess whether genotype PC correction offers additive benefit over SVA alone across populations with differing levels of stratification.

**Figure 2.**
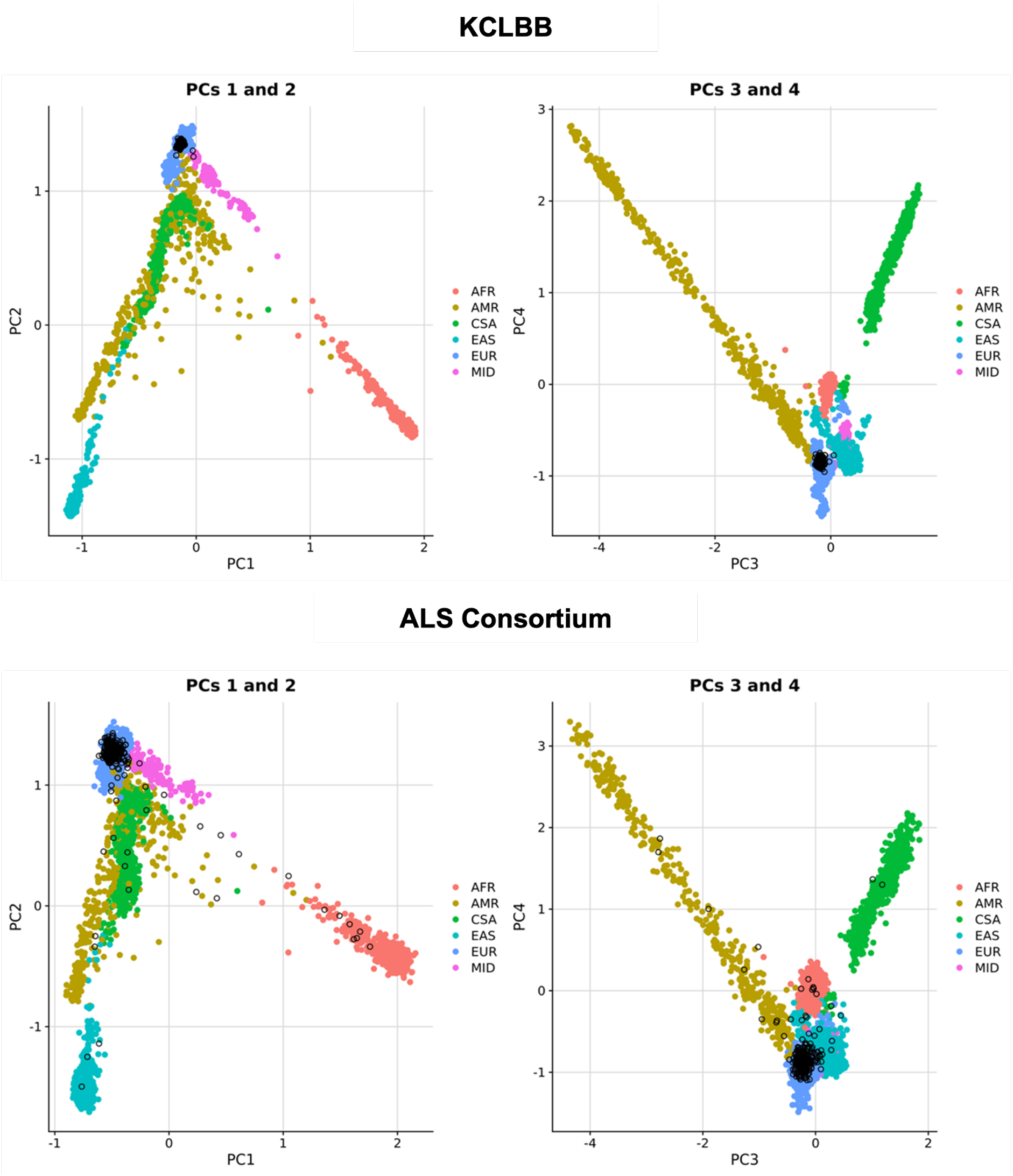
Global ancestry projections of study cohorts onto reference populations. PCA demonstrating the genetic ancestry of the KCLBB (top) and ALS Consortium (bottom) cohorts. Target samples (black open circles) are projected onto a space defined by the first four PCs of a diverse global reference panel (coloured points). Reference populations include African (AFR) in red, American (AMR) in yellow, Central/South Asian (CSA) in green, East Asian (EAS) in teal, European (EUR) in blue and Middle Eastern (MID) in pink. The first two PCs characterise continental level genetic variation, while subsequent components show finer sub-continental population structure.

### Differential Expression Analysis Across Modelling Frameworks

*Summary statistics per model.* Differential expression results diverged substantially between cohorts as model complexity increased (Table 1). In KCLBB (n = 26,008 genes), the Naïve model identified the largest number of DEGs (3,052; including 14 known ALS genes), with progressive latent variable correction resulting in a stepwise reduction in total discoveries. The SV-only model was most conservative (2,350 DEGs; 11 known ALS genes), while the SV+PC model partially recovered signal, identifying 2,443 DEGs and 12 known ALS genes, notably recovering *SCFD1* absent from the SVA-only model.

**Table 1.**
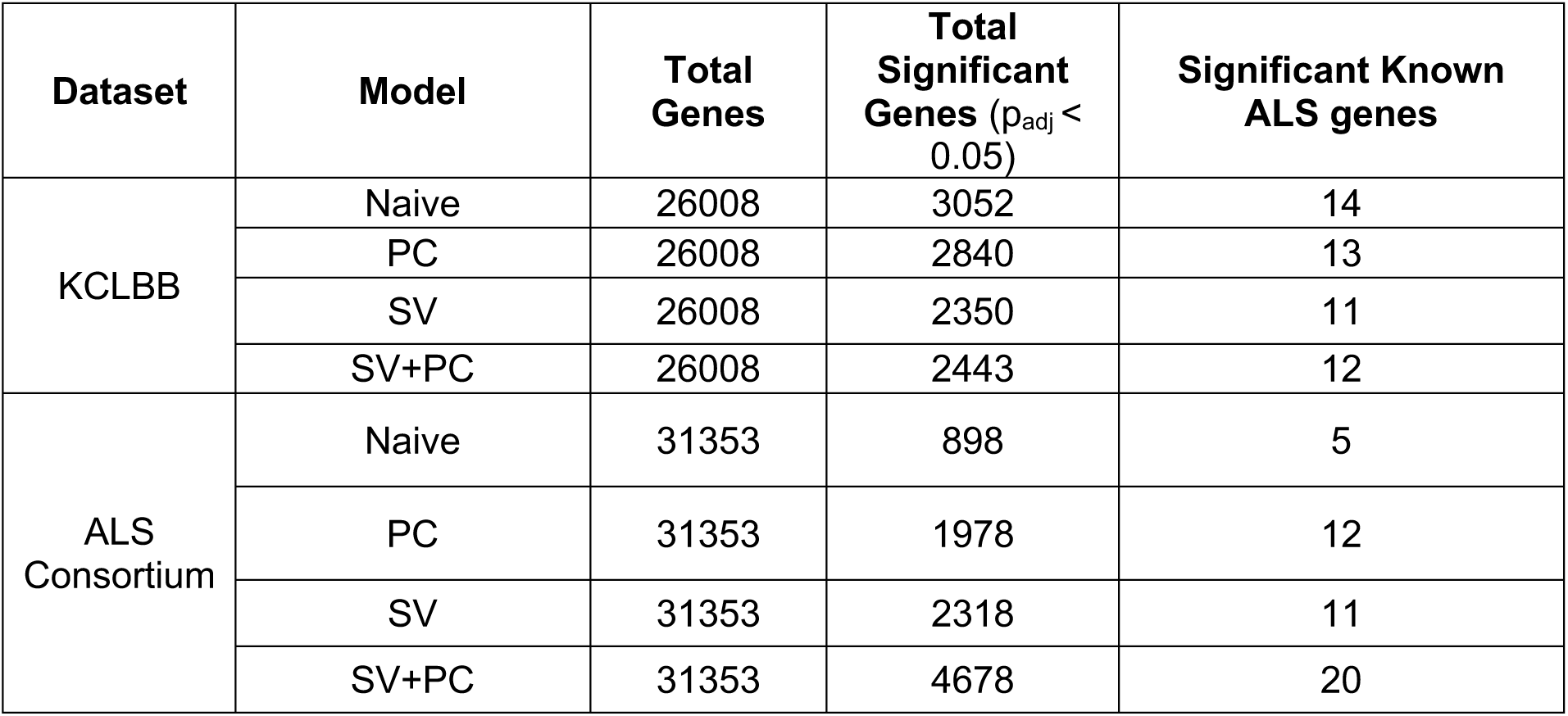
Summary Statistics for Differential Expression Modelling Frameworks.

In contrast, ALS Consortium (n = 31,353 genes) showed a consistent increase in discovery power with model complexity. The Naïve model identified only 898 DEGs and five known ALS genes, failing to recover any high-confidence Mendelian genes. The PC model more than doubled both total discoveries (1,978 DEGs) and ALS gene recovery (12 genes). The SV model further increased total DEGs (2,318) but marginally reduced ALS-specific recall (11 genes). The SV+PC framework was the most sensitive, identifying 4,678 significant genes and 20 known ALS genes, nearly double the SV-only model, retaining every locus identified across all simpler frameworks (Figure 3).

**Figure 3.**
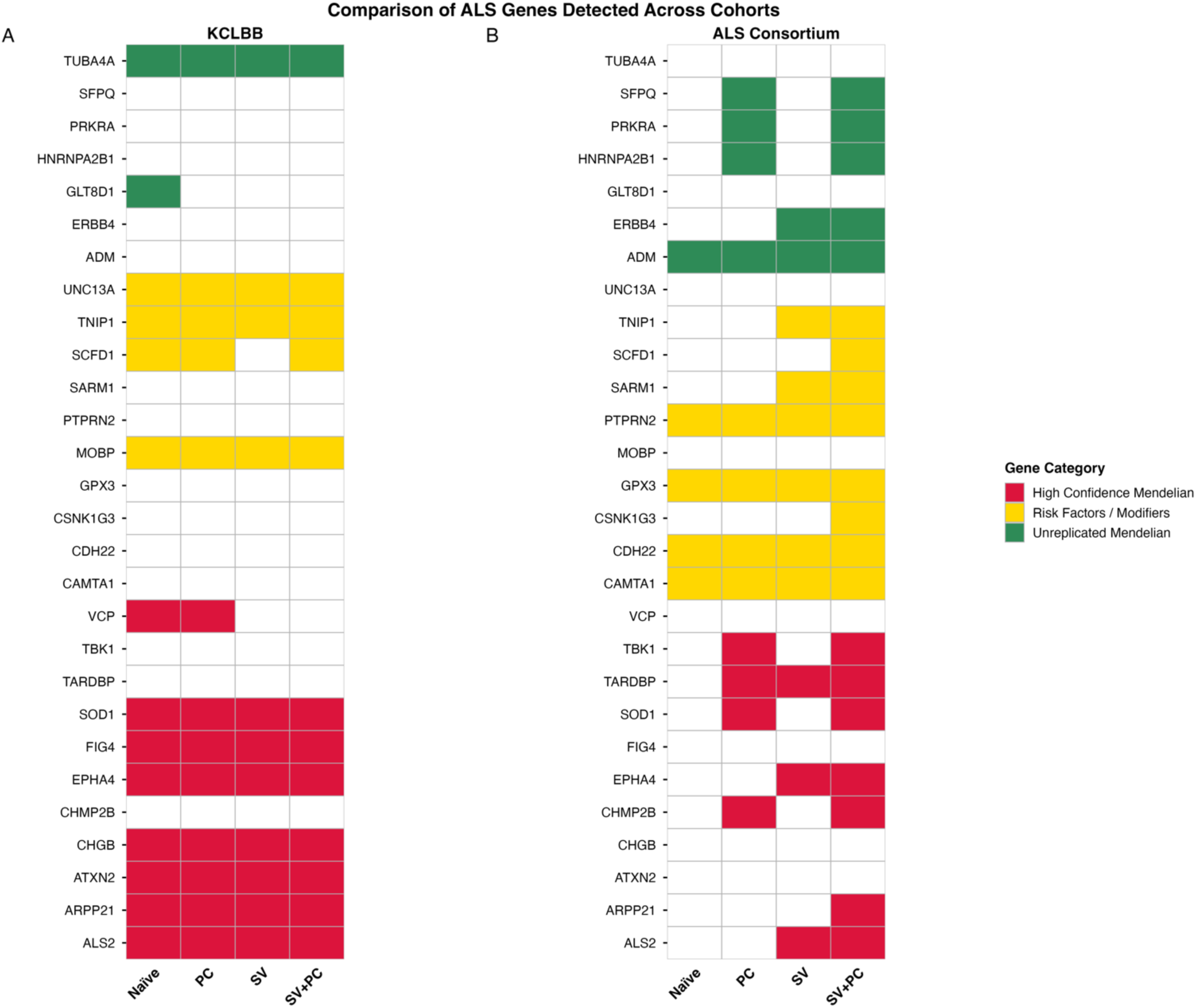
Comparative sensitivity of differential expression frameworks in recovering known ALS genes. This heatmap illustrates the detection of the curated set of known ALS genes across the four statistical modelling frameworks in KCLBB and ALS Consortium cohorts. Genes are categorised by evidence level: High Confidence Mendelian (red), Risk Factors/Modifiers (yellow), and Unreplicated Mendelian (green).

*Overlap of significant genes.* Intersection analysis revealed a high core consensus in KCLBB, with 2,114 genes shared across all four models, whereas ALS Consortium showed substantially lower consensus (590 genes shared). The SV+PC model identified the most unique discoveries in ALS Consortium (n = 1,691), and SV-based models jointly captured 1,133 genes absent from both the Naïve and PC-only frameworks, indicating that transcriptomic correction is prerequisite to unmasking this signal in the more heterogenous cohort (Figure 4).

**Figure 4.**
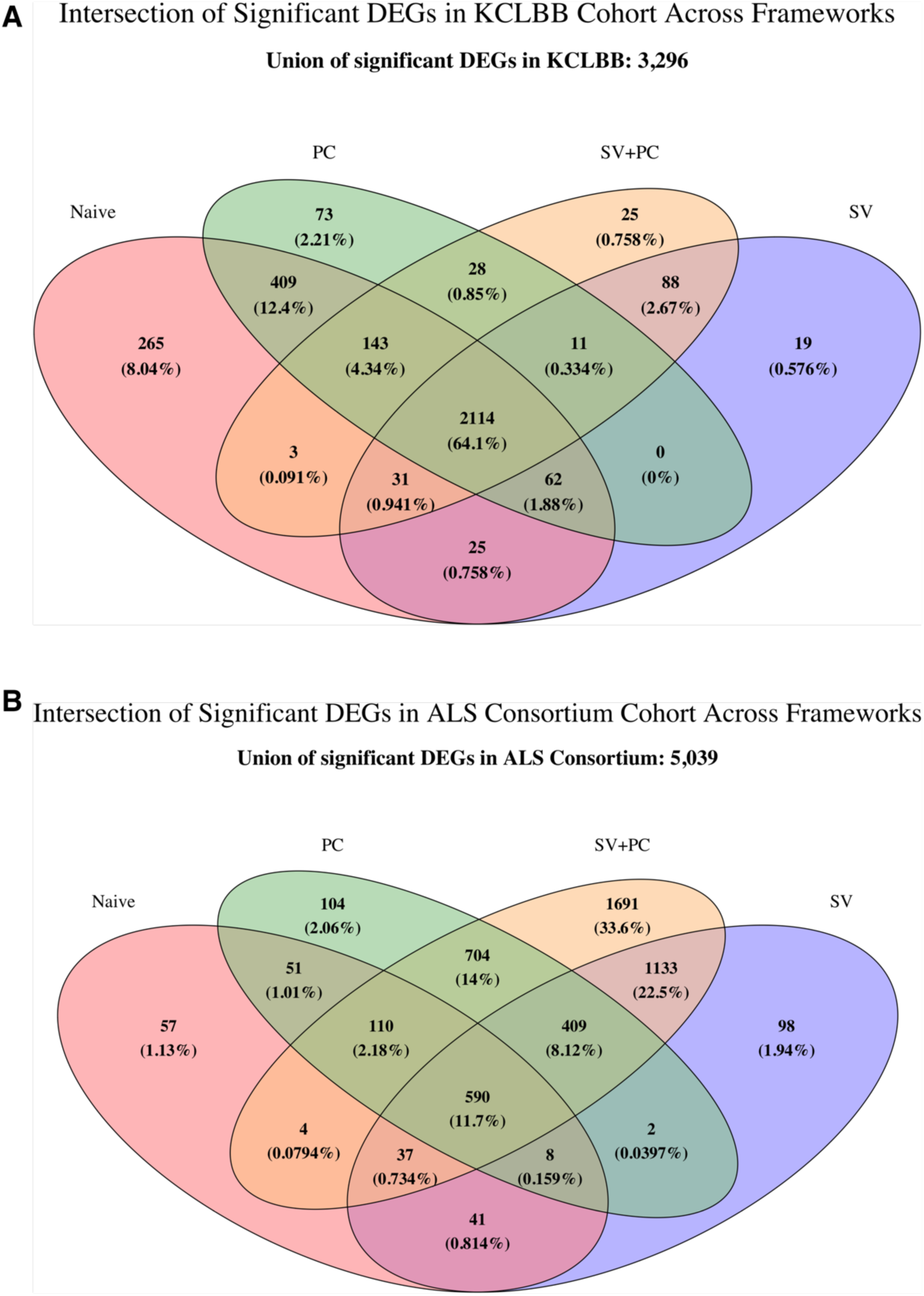
Comparative overlap of significant genes across modelling frameworks. Venn Diagrams illustrating the intersection of significant DEGs (p_adj_ < 0.05) for the KCLBB (A) and ALS Consortium (B) cohorts. Analysis was performed across the four frameworks: Naïve (red), PC (green), SV (blue) and SV+PC (yellow).

*Intra-dataset effect size concordance.* LFC estimates remained highly stable across all frameworks in KCLBB (minimum Pearson *r* = 0.941 across all filtered genes), while ALS Consortium showed greater sensitivity to model choice, with Pearson’s correlation dropping to *r* = 0.803 between the Naïve and SV+PC frameworks across the full transcriptome. Critically, in both cohorts the SV vs SV+PC comparison yielded the highest concordance, indicating that PC inclusion preserves SV-derived effect sizes magnitudes and rankings while expanding discovery (full statistics in Supplementary Table S3).

### Cross-Dataset Consistency and Replicability

*Effect size concordance across cohorts.* Cross-dataset correlation analyses revealed a consistent and progressive improvement in effect size consistency with increasing model complexity (Figure 5; Table 2). For the union of significant genes, the Naïve model yielded poor cross-cohort concordance (Pearson *r* = 0.228; Spearman ρ = 0.326), reflecting the dominance of dataset-specific technical noise in uncorrected models. The addition of PCs alone provided only marginal improvement (*r* = 0.303; ρ = 0.406), while SV correction substantially enhanced concordance (*r* = 0.541; ρ = 0.688). The SV+PC framework achieved the highest cross-dataset agreement across 5,961 genes (*r* = 0.572), with bootstrap resampling confirming significant improvement in consistency of effect size magnitude over the SVA-only model (Δ*r*: 0.027; 95% *CI* [0.004, 0.055]; see Supplementary Figure S2). However, Spearman rank correlation displayed a significant decrease between SV and SV+PC models (Δρ: −0.016; 95% *CI* [−0.020, −0.012]; see Supplementary Figure S2), suggesting that PC correction refines effect size magnitudes without fully preserving gene rankings across cohorts.

**Figure 5.**
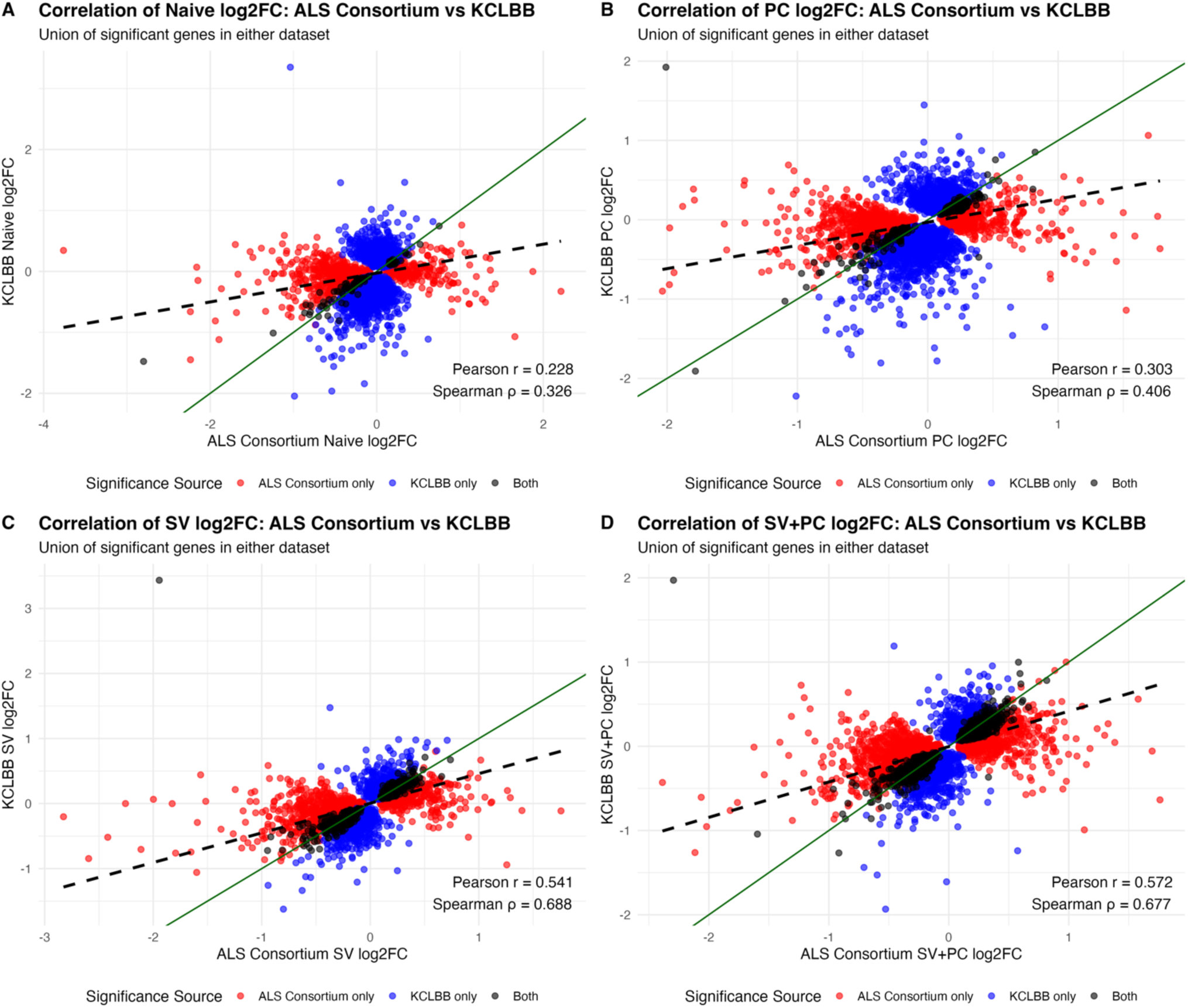
Correlation plots of log2 Fold Change (LFC) effect sizes for the union of significant genes between ALS Consortium and KCLBB across correction frameworks. Scatter plots depict the cross-dataset agreement under each statistical model: Naïve, PC, SV and SV+PC. The solid green line represents parity (y=x), while the dashed black line represents the linear regression fit. Points are colour-coded by significance source (Grey: both datasets; Blue: KCLBB only; Red: ALS Consortium).

**Table 2.**
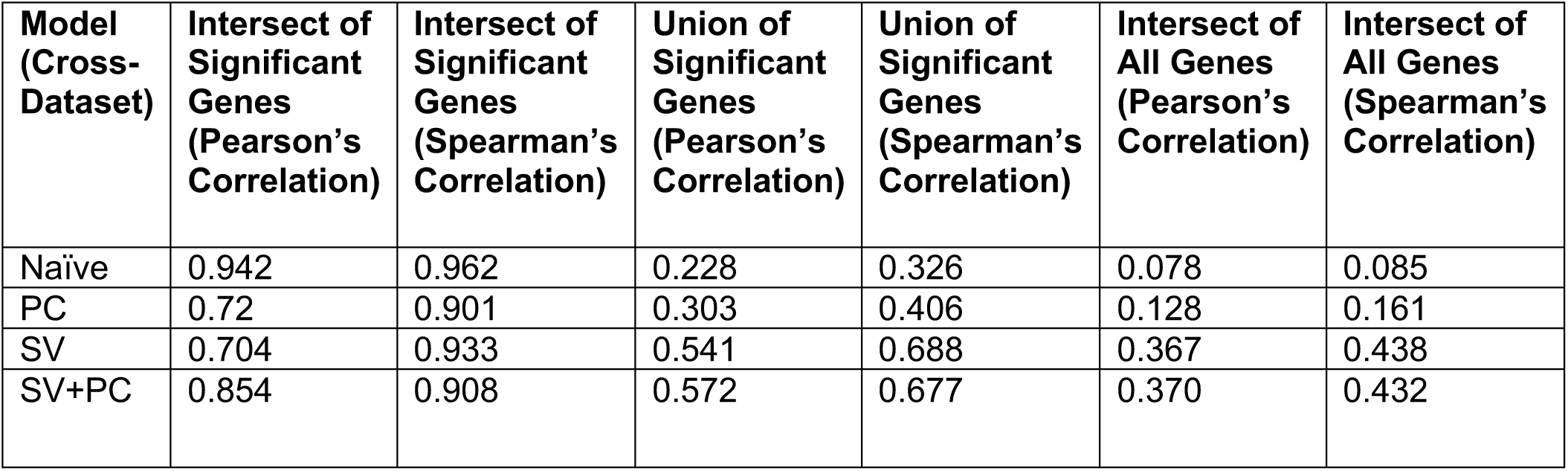
Cross-dataset correlation of effect sizes between ALS Consortium and KCLBB. Pearson (r) and Spearman (p) correlation coefficients were calculated to assess the stability and harmonization of the ALS transcriptomic signal between independent datasets. Results are reported for three distinct gene set: (i) the intersection of significant genes, representing the core consensus signal (padj < 0.05 in both cohorts); the union of significant genes, representing the model-dependent significant signal (padj < 0.05 in at least one cohort); (iii) the intersection of all genes, providing a global assessment of effect size stability across the entire filtered transcriptome (n = 25,895).

| Model (Cross-Dataset) | Intersect of Significant Genes (Pearson's Correlation) | Intersect of Significant Genes (Spearman's Correlation) | Union of Significant Genes (Pearson's Correlation) | Union of Significant Genes (Spearman's Correlation) | Intersect of All Genes (Pearson's Correlation) | Intersect of All Genes (Spearman's Correlation) |
| --- | --- | --- | --- | --- | --- | --- |
| Naïve | 0.942 | 0.962 | 0.228 | 0.326 | 0.078 | 0.085 |
| PC | 0.72 | 0.901 | 0.303 | 0.406 | 0.128 | 0.161 |
| SV | 0.704 | 0.933 | 0.541 | 0.688 | 0.367 | 0.438 |
| SV+PC | 0.854 | 0.908 | 0.572 | 0.677 | 0.370 | 0.432 |

This trend was consistent across all three gene sets. The core consensus signal, which are genes significant in both cohorts, expanded from just 88 genes in the Naïve to 1,160 genes in the SV+PC model. While the Naïve model showed the highest Pearson correlation for its intersection set (*r* = 0.942), this reflects a highly restricted and non-representative gene set. The SV+PC model maintained strong stability (*r* = 0.854; ρ = 0.908) over a substantially larger and more biologically representative shared set. At the global transcriptome level (25,895 genes), concordance improved from *r* = 0.078 in the Naïve model to *r* = 0.370 in the SV+PC framework, with the SV model achieving comparable global correlation (*r* = 0.367), indicating that SV correction drives the majority of global harmonisation.

*Replicability Index.* The Jaccard Similarity Index further supported the superiority of the SV+PC framework in recovering a shared biological signal (Table 3). The Naïve model demonstrated negligible overlap between cohorts ( *J* = 2.28%) despite identifying 3,862 significant genes in total. The PC model provided minimal improvement ( *J* = 6.29%), whereas SV correction yielded a substantial gain ( *J* = 17.3%). The SV+PC model achieved the highest replicability ( *J* = 19.5%), representing a nearly ten-fold increase over the uncorrected model. Bootstrap resampling confirmed that the 2.1% improvement in Jaccard Index conferred by adding PCs to the SV model was statistically significant (95% CI for Δ*Jaccard*: [0.012, 0.032], excluding zero; see Supplementary Figure S3). Among overlapping significant genes, directional concordance exceeded 99% across all models, indicating the increase in replicability was accompanied by consistency in direction of effect.

**Table 3.** Cross-dataset replicability and directional concordance of significant DEGs between KCLBB and ALS Consortium. For each model, the number of overlapping significant DEGs (intersection), the union of significant DEGs across both cohorts, the replicability of DEGs measured by the Jaccard Similarity Index and the proportion of overlapping significant DEGs with concordance direction of effect are reported.

| Model | Number of Overlapping Genes | Union of Significant Genes | Jaccard's Similarity Index (%) | Concordance of Effect Direction (%) |
| --- | --- | --- | --- | --- |
| Naive | 88 | 3862 | 2.28 | 100 |
| PC | 285 | 4533 | 6.29 | 99.9 |
| SV | 689 | 3979 | 17.3 | 99.3 |
| SV+PC | 1160 | 5961 | 19.5 | 99.7 |

### Biological Recall

Biological recall against a curated reference set of 66 evaluable known ALS genes (Supplementary Table S1) demonstrated a stepwise improvement in model sensitivity with increasing complexity (Table 4). The Naïve model failed to identify any known ALS genes in both cohorts simultaneously (Recall = 0.0%; Precision = 0.0%;), demonstrating that uncorrected latent variation obscures disease-relevant signals. The PC model achieved marginal improvement, recovering only *SOD1* (Recall = 1.52%; Precision = 0.351%). The SV model identified three consensus genes: *ALS2*, *EPHA4*, *TNIP1*; increasing recall and precision to 4.55% and 0.435% respectively. The SV+PC framework achieved the highest biological recall (9.10%) and ‘apparent’ biological precision (0.517%), recovering six known ALS genes across both cohorts: *ALS2*, *SCFD1*, *EPHA4*, *SOD1*, *TNIP1* and *ARPP21*. Notably, *SCFD1* was uniquely recovered by the SV+PC model and absent from all simpler frameworks, indicating that the simultaneous integration of genomic and transcriptomic latent variables is required to unmask this signal.

**Table 4.**
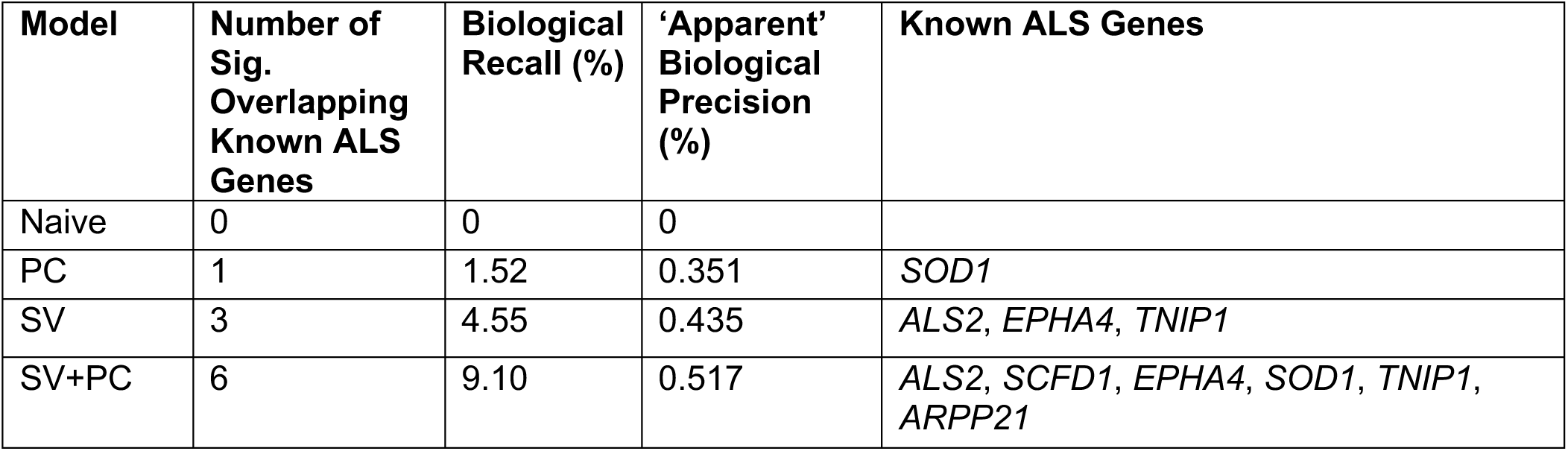
Biological recall and ‘apparent’ biological precision of known ALS risk genes across modelling frameworks.

| Model | Number of Sig. Overlapping Known ALS Genes | Biological Recall (%) | ‘Apparent’ Biological Precision (%) | Known ALS Genes |
| --- | --- | --- | --- | --- |
| Naive | 0 | 0 | 0 |  |
| PC | 1 | 1.52 | 0.351 | *SOD1* |
| SV | 3 | 4.55 | 0.435 | *ALS2*, *EPHA4*, *TNIP1* |
| SV+PC | 6 | 9.10 | 0.517 | *ALS2*, *SCFD1*, *EPHA4*, *SOD1*, *TNIP1*, *ARPP21* |

### Sensitivity Analysis

*Cross-dataset concordance.* Across all pairwise comparisons, the SV+PC model either significantly outperformed the SV model or showed no statistically significant difference in Pearson’s correlation (Figure 6A). ALS Consortium PC1 showed no significant improvement with any KCLBB PC interval except KCLBB PCs 1-10; ALS Consortium PCs 1-3 with KCLBB PCs 1-5 and 1-7; ALS Consortium PCs 1-7 with KCLBB PCs 1-5 and 1-7; and ALS Consortium PCs 1-10 with KCLBB PCs 1-5. All remaining combinations demonstrated a significant increase in effect size concordance for the SV+PC model relative to SV-only. In contrast for Spearman’s rank concordance, SV+PC predominantly displayed a statistically significant decrease relative to SV with only the KCLBB PCs 1-10 with ALS Consortium PC1, PCs 1-5 and PCs 1-7 showing no difference in gene rankings between models (Figure 6B).

**Figure 6.**
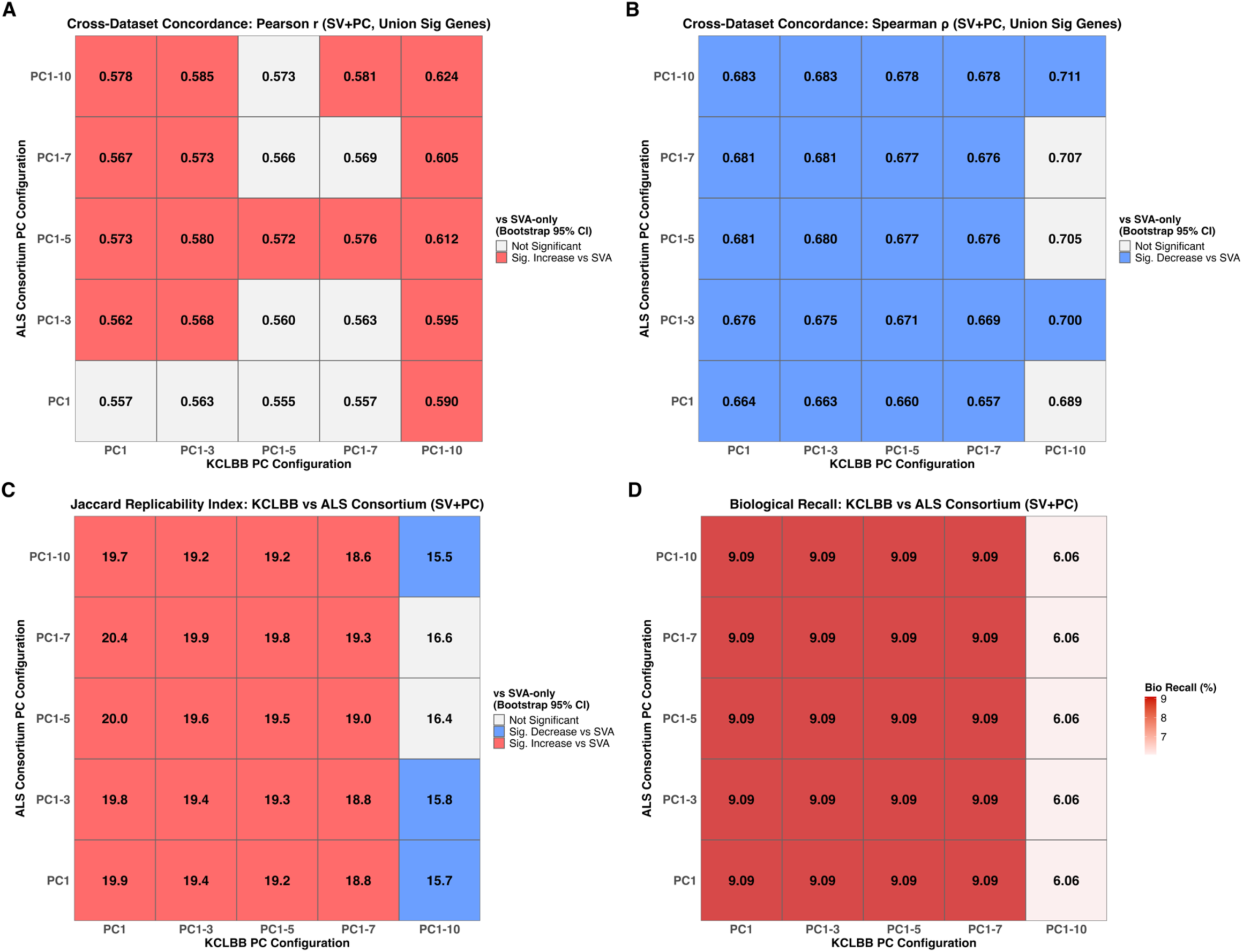
Sensitivity Analysis of SV+PC model performance across genotype PC intervals. Heatmaps display performance of the SV+PC model relative to SV-only across pairwise combinations of genotype PC intervals (PC1, PCs 1-3, PCs 1-5, PCs 1-7, PCs 1-10) for KCLBB and ALS Consortium. **(A)** Cross-dataset Pearson’s effect size concordance. **(B)** Cross-dataset Spearman’s concordance **(C)** Jaccard’s Replicability Index. For A-C, red cells denote a significant increase for SV+PC relative to SV, blue a significant decrease and white indicates no significant difference (bootstrap 95% CI). **(D)** Biological recall (3 significant figures); red intensity corresponds to higher recall.

*Replicability Index.* The SV+PC model significantly outperformed the SV model in most comparisons, with instability arising only at KCLBB PCs 1-10; this interval exhibited a statistically significant decrease when paired with ALS Consortium PC1, PCs 1-3 and PCs 1-10 and no statistically significant difference with ALS Consortium PCs 1-5 and PCs 1-7 (Figure 6C).

*Biological Recall.* The SV+PC model consistently achieved a higher recall than the SV model (4.55%), across all pairwise comparisons (Figure 6D). For all KCLBB PC intervals up to and including PCs 1-7, cross-dataset biological recall remained at 9.09% for all ALS Consortium configurations. However, at KCLBB PCs 1-10, biological recall decreased to 6.06% across all ALS Consortium PC combinations.

Overall, the results were consistent across the majority of PC interval combinations with instability arising primarily at KCLBB PCs 1-10.

## Discussion

This study provides the first systematic evaluation of combined genotype-derived PCs and expression-derived SV correction within a differential expression analysis framework, using two independent post-mortem ALS RNA-seq datasets. Across all evaluation metrics (cross-cohort replicability, biological recall and effect size concordance), the simultaneous inclusion of SVs and PCs consistently outperformed either correction strategy in isolation, revealing important insights into the performance of latent variable correction strategies.

Our intra-dataset analysis revealed a striking divergence in how model complexity affected the two cohorts. In the heterogenous ALS Consortium cohort, which also carried a severe case-control imbalance, increasing model complexity drove a substantial increase in sensitivity, with significant genes rising from 898 in the Naïve model to 4,678 in the SV+PC framework, and known ALS gene recovery increasing from 5 to 20. This suggests that in heterogenous populations, uncorrected latent variation actively suppresses genuine biological signal, and that dual correction helps to unmask it. In contrast, the homogenous KCLBB cohort showed a reduction in total discoveries as complexity increased, with a stable core of 2,114 genes retained across all models. Rather than indicating a loss of sensitivity, this pattern suggests that in genetically homogenous datasets, combined latent variable correction acts as a high-pass filter, pruning unreliable, noisy signals to refine the true biological core without fundamentally altering underlying effect sizes. These contrasting patterns highlight that the benefit of PC correction is not limited to multi-ethnic cohorts but also in largely homogeneous populations, as PC correction provides a precision gain that partially mitigates the risk of losing biological signal by SV correction alone.

A key concern in combined latent variable modelling is redundancy, if SVs and PCs capture overlapping variance, their simultaneous inclusion risk instability or overcorrection. Our multicollinearity analyses directly addressed this, demonstrating no significant associations between SVs and PCs in either cohort. This orthogonality is reinforced by the pattern of cross-dataset results, with SVs driving the largest single improvement in cross-cohort replicability, while the addition of PCs contributed a further independent and statistically significant gain in Jaccard index (Δ*Jaccard* = 2.1%; 95% *CI* [0.012, 0.032]). This additive structure is precisely what would be expected if SVs and PCs are addressing non-overlapping sources of variation: transcriptomic technical noise and genomic population structure, respectively. This is further illustrated by the association of ALS Consortium PC1 with disease status, which rather than representing a confounding variable, likely reflects population stratification covarying with disease susceptibility in a cohort with a marked case control imbalance. PC1 effectively partitions population structure from primary disease signal, consistent with the substantial gain in sensitivity observed in ALS Consortium under the SV+PC framework. This supports the broader principle that PCs can capture population-stratified variation, which is associated with status, a source of confounding that SVs alone, operating exclusively on expression data, is not constrained to resolve.

This distinction has practical implications for model design. Prior ALS transcriptomic studies which predominantly employ SV-based correction [14] or expression PC correction [15] in isolation, while other studies have incorporated genotype PCs primarily as stratification control in expression quantitative trait loci (eQTL) analyses [30]. Our findings suggest that these approaches, taken independently, are suboptimal. As the low cross-dataset replicability of the PC-only model (*Jaccard* = 6.29%) confirms that genotype-based correction alone is inadequate in addressing transcriptomic confounding, while the near-equivalent correlations of the SV and SV+PC models across gene sets suggest that SVs drives the primary harmonisation of the data structure, but PCs then provide a fine-tuning effect, recovering biologically relevant signals (including known ALS genes absent from the SV-only framework) which remain trapped beneath a residual layer of population stratification. Notably, bootstrap resampling confirmed that this fine-tuning effect is specific to effect size magnitude, as addition of PCs produced a statistically significant increase in Pearson correlation but a significant decrease in Spearman rank correlation between cohorts. This divergence may reflect the compression of LFC values towards zero following PC addition, which improves the overall linear relationship between cohorts but increases rank instability among the dense gene cluster with modest effect sizes, where trivial differences in magnitude determines rank order; a trade-off which warrants consideration when rank-based interpretation is prioritised. These findings suggest that differential expression analysis studies relying on a single correction approach, whether SV or PC correction alone, may have missed signals and report findings with inflated dataset-specific noise, a limitation that the hybrid framework directly addresses.

The cross-dataset findings represent the central contribution of this study. The progressive improvement in concordance across all metrics as model complexity increases demonstrates that dual correction is a statistical refinement which improves the recovery of a reproducible ALS transcriptomic signature. The apparent high correlation of the Naïve model’s intersection gene set (*r* = 0.942) is an artefact of selection bias, reflecting a tiny subset of 88 genes with extreme signals that survived dataset-specific noise, as evidenced by a replicability index of just 2.28%. Beyond this, increasing model complexity produced consistent and simultaneous gains in both size of the shared DEG set and the consistency of effect sizes, with the SV+PC framework reaching 1,160 shared genes and a Pearson correlation of 0.572 for the union of significant genes, compared to just 88 genes and *r* = 0.228 in the Naïve model. The expansion of shared biology without sacrificing effect size stability and direction is the defining characteristic of the combined model and distinguishes it from single-correction approaches, which each sacrifice either sensitivity or precision.

Biological recall and ‘apparent’ precision against a curated reference set of 66 established known ALS genes further validated this conclusion. The Naïve model failed to recover any known ALS genes across both cohorts, and the PC-only model recovered only *SOD1*. The SV+PC framework doubled the recall of the SV-only model (9.10% vs 4.55%), uniquely recovering *SOD1*, *SCFD1* and *ARPP21*, which suggests that these are all genes obscured by the residual population structure and technical confounding that single-correction strategies fail to resolve. This indicates that for genes with subtle or context-dependent transcriptional effects, the dual-correction framework provides the statistical precision necessary to cross the threshold of significance in both independent cohorts simultaneously. The marginal gain in ‘apparent’ precision from SV-only to SV+PC further supports that notion that dual correction prunes residual noise to refine the biological signal rather than inflating the significant gene set.

Sensitivity analysis indicated that the performance of the SV+PC framework is broadly robust to the number of genotype PCs included, as it either significantly outperformed or was statistically indistinguishable from the SV-only model across Pearson’s concordance, Jaccard replicability and biological recall across most of the 25 pairwise combinations. Pearson’s concordance never significantly decreased and biological recall remained consistently higher than SVA across all configurations, suggesting that effect size and biological signal improvements are stable properties of the combined model. These findings imply that results are not specific to a given PC choice, which is reassuring for practical application of this framework. Instability arose predominantly at KCLBB PCs 1-10, where replicability reduced across ALS Consortium pairings and biological recall fell from 9.09% to 6.06%, though this remained above the SVA baseline of 4.55%, consistent with overadjustment at this interval. This is supported by the KCLBB scree plot, which suggests negligible variance explained beyond PC5, highlighting that it is unlikely to capture genuine population structure in this homogenous cohort. The persistent decrease in Spearman’s rank concordance across most configurations further corroborates that PC addition structurally influences gene rankings through effect size compression, largely independent to the number of PCs chosen. Collectively, the sensitivity analysis supports that PC number should be guided by the Scree plot elbow for this analysis. These findings strengths the central conclusion of this study by demonstrating that the superiority of the SV+PC framework is robust across a wide range of analytical choices.

While these findings presented here are specific to ALS post-mortem brain tissue, the methodological principle extends naturally to any transcriptomic study in which population stratification and technical heterogeneity co-exist. Differential expression analysis studies in other complex diseases are likely to encounter the same dual-layer confounding characterised here, and the SV+PC framework represents a generalisable solution to this problem.

Despite the improvements demonstrated by the SV+PC framework, the overall biological recall of 9.10% highlights the inherent challenges of ALS transcriptomics. The majority of curated known ALS genes remain undetected, which likely reflects several compounding factors. Many genetic drivers of ALS may exert effects too subtle for the statistical power of the cohorts used, particularly given the case-control imbalances present in both datasets. Furthermore, a subset of known ALS genes may not act through altered steady-state mRNA levels, but through protein structure/function [31] or post-translational mechanisms [32] invisible to bulk differential expression analysis. Capturing these signals will likely require larger meta-analyses or complementary approaches such as splicing quantitative trait loci. Additionally, the consistently low ‘apparent’ precision across all models suggests that some of the replicated transcriptomic signal may reflect dysregulated biology not yet catalogued in the known ALS gene reference set.

A further limitation is the reliance on bulk RNA-seq, which averages transcriptional signal across heterogenous cell populations. Cell-type specific signals, such as those exclusive to motor neurons or specific glial populations, are likely diluted in whole-tissue samples [33], reducing the sensitivity to detect disease-relevant expression changes in the cell types most affected in ALS. Future work integrating single-cell RNA-seq data would be valuable for resolving these cell-type specific effects.

The accuracy of PC-based ancestry inference in the ALS Consortium was also constrained by a technical feature of the WGS-derived genotype data. Homozygous reference genotypes were encoded as “./.” and were therefore indistinguishable from genuinely missing calls. Consequently, standard missingness based QC excluded a large proportion of variants, reducing the genomic coverage available for PCA. If anything, this would be expected to reduce the accuracy of ancestry inference by limiting PC computation to a small subset of markers. Nevertheless, the reference-projected ancestry clusters in ALS Consortium appeared comparably well resolved to those in KCLBB, which had denser genome coverage, suggested that the retained variants still captured the major axes of population structure.

Both cohorts also suffer from limited ancestral diversity, with KCLBB representing an almost entirely European population and ALS Consortium containing only modest non-European representation. This restricts the generalisability of the shared transcriptomic signature across global populations and means that the full benefit of PC correction in genuinely diverse cohorts remains to be characterised. Encouragingly, however, the SV+PC framework demonstrated superiority over all simpler models even in these conditions, suggesting that the approach is likely to perform at least as well in more ancestrally diverse datasets. Future studies incorporating cohorts with greater representation across ancestral groups will be necessary to fully validate this.

## Conclusion

This study provides the first systematic evaluation of combined genotype PC and expression-derived SV correction within a differential expression analysis framework. We demonstrated that the transcriptomic landscape of complex diseases like ALS, is obscured by latent technical and population-specific variation, and that traditional single-correction approaches are less able to recover reproducible disease signals across independent datasets. The hybrid SV+PC model consistently outperformed simpler models across all evaluation metrics, achieving the highest cross-dataset replicability, biological recall and effect size consistency in both homogenous and heterogeneous datasets. Critically, sensitivity analysis suggested these findings are relatively robust to the number of PCs included, with performance remaining stable across most PC interval combinations, and instability arising predominantly beyond the Scree plot elbow for KCLBB, consistent with overadjustment in a homogeneous population. The mechanistic basis of this improvement lies in the conceptually distinct variance structures targeted by each correction layer: SVs resolves dominant transcriptomic noise such as batch effects and unmeasured biological heterogeneity, while genotype PCs are theoretically positioned to address expression differences driven by population structure that expression-based methods alone cannot fully resolve. However, since we cannot determine the extent to which SVs independently capture population stratification, or whether PC association with site partly reflect technical batch effects genotype data collection, the precise mechanistic contribution of each correction layer remains difficult to discern. These findings have direct implications beyond ALS. Any transcriptomic study is likely to benefit from this dual-correction approach. We recommend the routine incorporation of genotype PCs alongside expression SVs as a standard practice in differential expression analysis study design, and propose that future work investigates whether equivalent gains are achievable using RNA-derived markers for population structure in studies lacking matched genotype data.

## Contributions

YA, OP and AI conceived the study; YA performed the analyses, generated the first article draft, figures and tables; AI and OP supervised the study; NYGC ALS Consortium provided the bulk RNA-seq and WGS data; JJ, AA, ST, AI and OP, provided feedback throughout the course of the work; YA, OP and AI led the writing of the final version of the manuscript; YA, JJ, AA, ST, OP, AI, reviewed and approved the final manuscript.

## Supporting information

List of NYGC ALS Consortium Principal Investigators and Sites

## Acknowledgements

We would like to thank the London Neurodegenerative Diseases Brain Bank and the NYGC ALS Consortium for generating and maintaining the transcriptomic datasets. All NYGC ALS Consortium activities are supported by the ALS Association (ALSA, 19-SI-459) and the Tow Foundation, as well as The Target ALS Human Postmortem Tissue Core, New York Genome Center for Genomics of Neurodegenerative Disease, Amyotrophic Lateral Sclerosis Association and TOW Foundation. We also acknowledge the relevant funding bodies that support ongoing ALS research, including the NIHR Maudsley Biomedical Research Centre and collaborating international organisations. We thank people with MND and their families for their participation in the initiatives that led to the generation of the data used in this project. The authors acknowledge use of the King’s Computational Research, Engineering and Technology Environment (CREATE) (https://create.kcl.ac.uk), which is delivered in partnership with the National Institute for Health and Care Research (NIHR) Biomedical Research Centres at South London and Maudsley and Guy’s and St. Thomas’ NHS Foundation Trusts and part-funded by capital equipment grants from the Maudsley Charity (award 980) and Guy’s and St.Thomas’ Charity (TR130505).

## Declarations

OP provides consultancy services for UCB Pharma. Post-mortem tissue samples from King’s College London were collected under the ethical approval of the MRC London Neurodegenerative Diseases Brain Bank and under the regulations of the Human Tissue Act UK 2014. All post-mortem tissue was donated to the MRC London Neurodegenerative Diseases Brain Bank under standard ethical and Human Tissue Act procedures, with informed consent provided by the next of kin. Data generated from this material were anonymized and analysed on a high-performance computing cloud (https://www.maudsleybrc.nihr.ac.uk/facilities/rosalind/) with data protection protocols in accordance with Department of Health Policy (UK) and the security standards set by the National Data Guardian. Ethical approval to process and analyse post-mortem samples stored at King’s College London was provided by a local ethics committee at the Institute of Psychiatry, Psychology & Neuroscience, King’s College London, and the MRC London Neurodegenerative Diseases Brain Bank. The NYGC ALS Consortium samples presented in this work were acquired through various institutional review board (IRB) protocols from member sites and the Target ALS postmortem tissue core and transferred to the NYGC in accordance with all applicable foreign, domestic, federal, state, and local laws and regulations for processing, sequencing, and analysis. The Biomedical Research Alliance of New York (BRANY) IRB serves as the central ethics oversight body for NYGC ALS Consortium. Ethical approval was given. Informed consent has been obtained from all participants

## Availability of Data and Materials

The KCL BrainBank datasets are available upon reasonable request from the corresponding author. The NYGC ALS Consortium dataset is available upon approval by the NYGC ALS Consortium. All scripts are available at https://github.com/yadusayanappulingam/Evaluating-SVs-and-PCs-in-DEA.

