## Supplementary Methods and Figures for "Exploring transcriptomic and genomic latent variable correction approaches in differential expression analysis"

### Multi-collinearity and Covariate Selection

Due to the inclusion of multiple latent variables derived from genotype and phenotype data, there is a risk of multicollinearity in the linear model. This can lead to unstable estimates of fold change and reduced statistical power during our differential expression analysis when combining SVs and PCs into a singular model. To evaluate potential multi-collinearity, systematic association testing was conducted to assess the independence of latent factors from measured clinical and technical covariates, as well as the internal orthogonality of latent variables within the modelling framework. Pearson's correlation coefficients were utilised to assess relationships between continuous variables, whilst independent t-tests and One-way Analysis of Variance (ANOVA) was employed to evaluate associations involving binary and multi-level categorical variables respectively.

### Model Specifications

1. **Naïve model**:
   The baseline model which only includes primary clinical and technical covariates.

$$Naive_{KCLBB} \sim Sex + Age_{Centred} + RIN_{Cat} + PMD_{Centred} + Status$$

$$Naive_{ALS\ Consortium} \sim Sex + Age_{Centred} + RIN_{Cat} + Site + Status$$

   In the KCLBB model, PMD was included whereas in the ALS Consortium model, PMD was excluded due to high missingness. Additionally, Site was added to account for the different sites used for sample collection in the ALS Consortium dataset. KCLBB did not require this covariate as all samples were from the same collection site.

2. **PC Model**:
   This model adds genotype-derived PCs to the Naïve model.

$$PC_{KCLBB} \sim Sex + Age_{Centred} + RIN_{Cat} + PMD_{Centred} + PC1 + PC2 + PC3 + PC4 + PC5 + Status$$

$$PC_{ALS\ Consortium} \sim Sex + Age_{Centred} + RIN_{Cat} + Site + PC1 + PC2 + PC3 + PC4 + PC5 + Status$$

3. **SV Model:**
   This model adds expression derived SVs to the Naïve model.

$$SV_{KCLBB} \sim Sex + Age_{Centred} + RIN_{Cat} + PMD_{Centred} + SV1 + Status$$

$$SV_{ALS\ Consortium} \sim Sex + Age_{Centred} + RIN_{Cat} + Site + SV1 + SV2 + SV3 + Status$$

4. **SV+PC Model (Combined):**
   This final model includes both genetic PCs and expression SVs to evaluate the additive benefit of accounting for both technical expression heterogeneity and population structure simultaneously.

$$SV + PC_{KCLBB} \sim Sex + Age_{Centred} + RIN_{Cat} + PMD_{Centred} + SV1 + PC1 + PC2 + PC3 + PC4 + PC5 + Status$$

$$SV + PC_{ALS\ Consortium} \sim Sex + Age_{Centred} + RIN_{Cat} + Site + SV1 + SV2 + SV3 + PC1 + PC2 + PC3 + PC4 + PC5 + Status$$

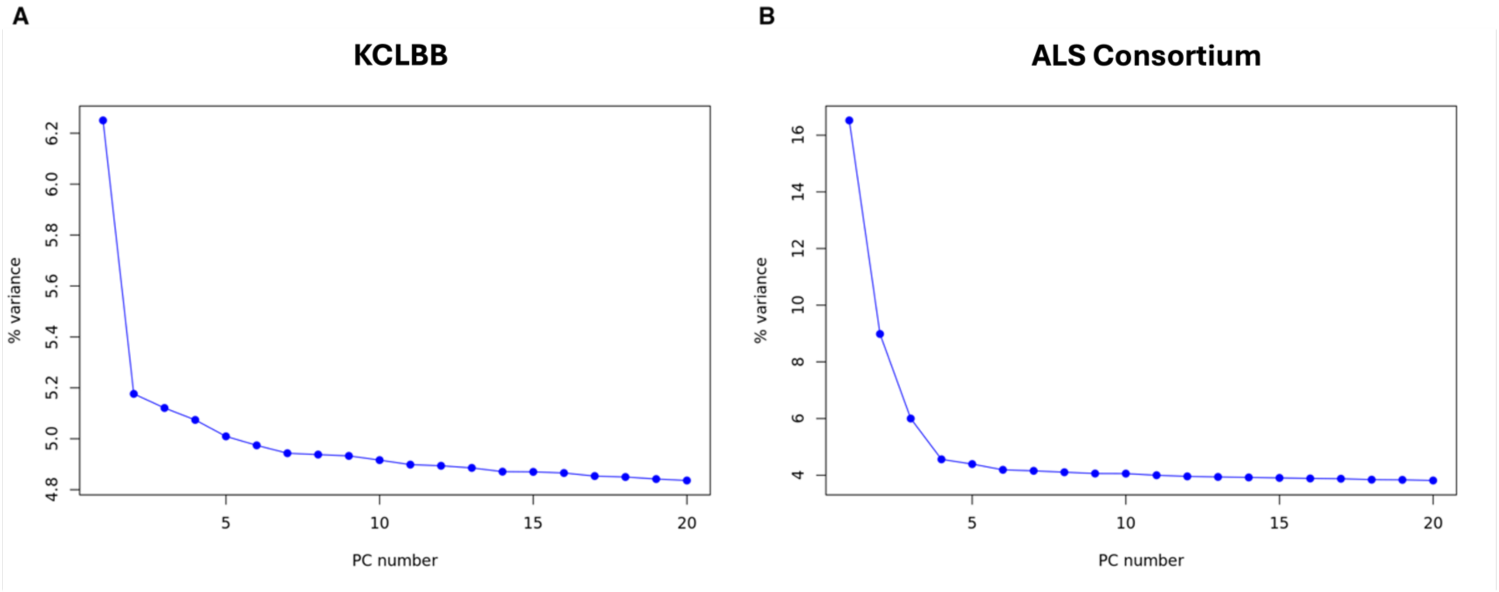


**Supplementary Figure S1. Principal component analysis scree plots for KCLBB (A) and ALS Consortium (B).** The y-axis represents the proportion of total variance explained per PC and the x-axis denotes the number of PCs.

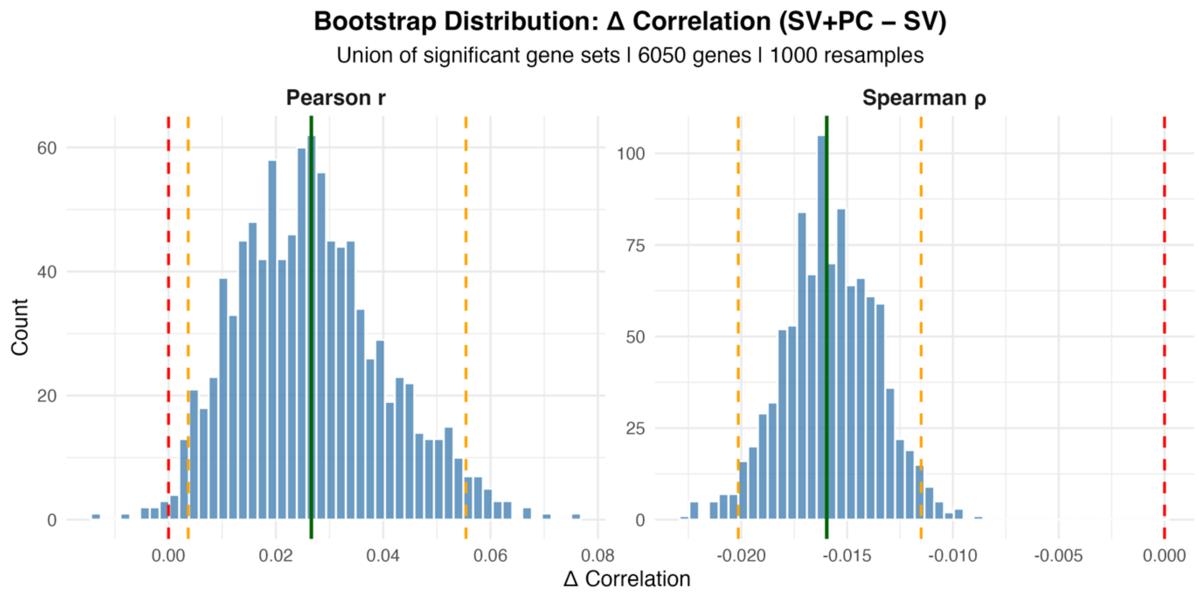


**Supplementary Figure S2. Bootstrap distribution of the difference in Pearson (left) and Spearman (right) correlation between SV-only and SV+PC models.** $\Delta r$ was calculated as $r\ (SV + PC) - r(SV)$ and $\Delta\rho$ was calculated as $\rho(SV + PC)$ - $\rho(SV)$ across 1000 bootstrap resamples of the union of significant genes across cross-dataset SV and SV+PC frameworks (6050 genes, resampled with replacement). The green vertical line represents the observed $\Delta Jaccard$. Orange dotted lines denote the 95% confidence interval. The red dashed line represents zero.

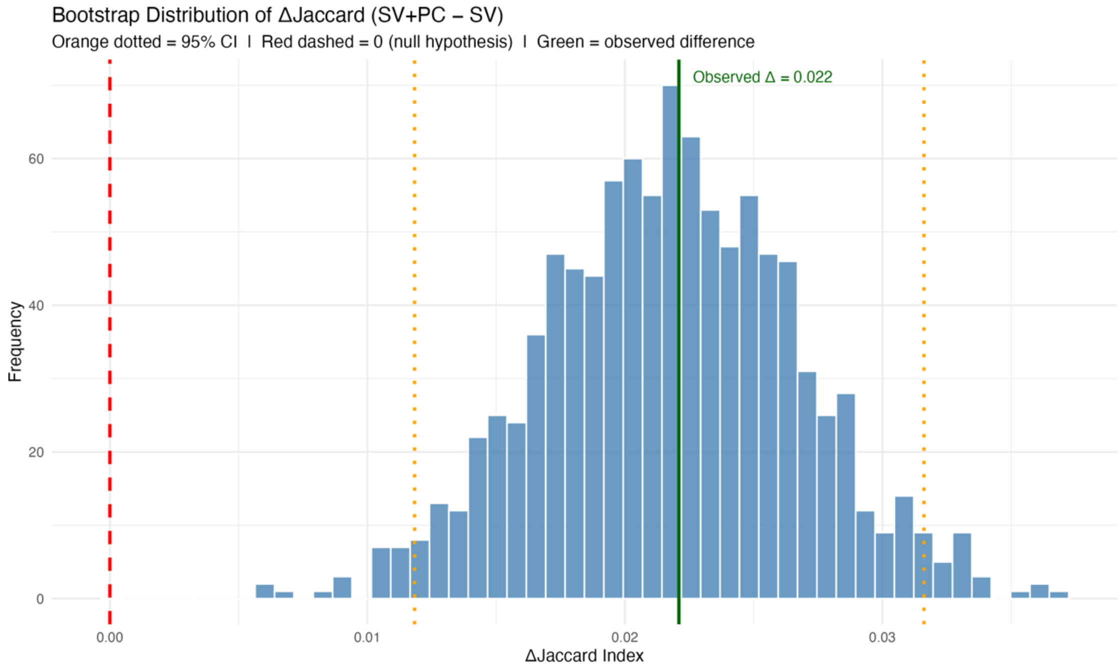


**Supplementary Figure S3. Bootstrap distribution of the difference in Jaccard Similarity Index between SV+PC and SV models.** $\Delta Jaccard$ was calculated as J($(SV) + PC$) – J($(SV)$) across 1000 bootstrap resamples of the shared gene universe (25985 genes, resampled with replacement). The green vertical line represents the observed $\Delta Jaccard$. Orange dotted lines denote the 95% confidence interval. The red dashed line represents zero.
